# How does individual variation in sociality influence fitness in prairie voles?

**DOI:** 10.1101/676858

**Authors:** Anne C. Sabol, Connor T. Lambert, Brian Keane, Nancy G. Solomon, Ben Dantzer

**Author notes:** Correspondence: Anne Sabol, 11200 SW 8^th^ St., Miami, FL 33199. Declarations of interest: none.

## Abstract

Comparative studies aid in our understanding of specific conditions favoring the initial evolution of different types of social behaviors, yet there is much unexplained intraspecific variation in the expression of social behavior that comparative studies have not yet addressed. The proximate causes of this individual variation in social behavior within a species have been examined in some species but its fitness consequences have been less frequently investigated. In this study, we quantified the fitness consequences of variation in the sociality of prairie voles (*Microtus ochrogaster*). We characterized sociality of voles in semi-natural enclosures using an automated behavioral tracking system paired with social network analyses to quantify the degree of spatial and temporal co-occurrence of different voles. We then assessed the relationship between sociality with mating success (number of different conspecifics with which an individual produced offspring) and reproductive success (total number of offspring surviving to first capture). We measured the number of social connections each individual had with all voles and only with opposite-sex voles by calculating unweighted degree through social network analyses. Both female and male voles varied in the number of social connections they had with all conspecifics and with opposite-sex conspecifics. Voles with an intermediate number of social connections with voles of both sexes had higher mating success overall. In our analyses that considered all social connections with voles of both sexes, voles with an intermediate number of social connections produced more offspring. Males with a very high or low number of social connections also had the lowest average body mass. Overall, our results suggest some limit on the fitness benefits of sociality. Although there was substantial individual-variation in our measure of vole social behavior, intermediate levels of social connections may be most favorable.

## Introduction

Sociality comes in many forms across a diversity of taxa, ranging from loose, temporary associations during breeding to long-term group associations whose members exhibit complex social behaviors such as cooperative hunting or alloparental care. Different forms of sociality are thought to provide a variety of fitness benefits, including shared resources, reduced predation, communication, or better access to mates (Eisenberg et al., 1972; Alexander, 1974; van Schaik 1983; Emlen, 1984; Krause and Ruxton, 2002, Silk, 2007). Sociality may also come with costs associated with group-living, including increased disease transmission (Ewald, 1994; Langwig et al., 2012; Kappeler et al., 2015), parasite load (Côté, and Poulinb, 1995; Whiteman and Parker, 2004), physiological stress (Creel et al., 2013), reproductive interference by other group members (Clutton-Brock et al., 1998; Lukas and Huchard, 2014), as well as the time and energy costs devoted to developing and maintaining social connections that could otherwise be devoted towards activities directly related to individual survival or reproduction such as foraging or parental care. Given that these fitness benefits and costs of sociality may vary in direction or magnitude due to ecological circumstances such as food abundance, weather patterns, or temporal variability in these factors (Emlen, 1994; Hatchwell and Komdeur, 2000; Shuster and Wade, 2003; Schradin and Pillay, 2005; Rubenstein and Lovette, 2007; Schradin et al., 2010; Shen et al., 2017), it is no surprise that there is considerable variation in the presence or expression of different types of social behaviors among species.

Comparative studies have been useful in revealing some of the factors favoring the evolution of different types of social behaviors. For example, phylogenetic comparative meta-analyses show that social monogamy is more commonly found in mammalian species that experience low population densities (Lukas and Clutton-Brock, 2013) and genetic monogamy is also more common in mammalian species with low population densities as well as those that exhibit parental care (Lambert et al., 2018). Other comparative studies have shown that cooperative breeding, where subordinate group members care for offspring produced by dominant breeders, is more likely to be found in mammalian species that live in arid areas (Faulkes et al., 1997; Lukas and Clutton-Brock, 2017) or in avian species that inhabit areas where rainfall is low and unpredictable (Jetz and Rubenstein, 2011). While these studies help illuminate some factors affecting the evolution of social behaviors like social monogamy or cooperative breeding, they typically cannot address the causes of intraspecific variation in sociality.

Sociality is not always expressed to the same level or in the same way within a species and there are often a variety of social strategies within the same species (Lott, 1991; Clutton-Brock, 1989; Getz et al., 1993). The magnitude of variation in the expression of social behavior within a species may not be as large as that among species but it requires explanation nonetheless. Although some studies have investigated some of the proximate causes of individual variation in specific social behaviors such as social monogamy (Young and Wang, 2004; Donaldson and Young, 2008; Ophir et al., 2008; Okhovat et al., 2015; Walum and Young, 2018) or alloparental care (Dantzer et al., 2017), fewer studies document its fitness consequences. This is not surprising as it requires detailed studies that document the social behavior of individuals and then relate their degree of sociality to their survival and reproduction. Most previous studies on this topic have been conducted in primates where investigators documented how the strength of social relationships (“social bonds”) impacted offspring survival, mating success, or longevity (e.g., Silk et al., 2003, 2009, 2010; Schülke, et al., 2010). By investigating the association between social behavior and fitness within a species, we may be able to understand how individuals balance the tradeoffs between the costs and benefits of sociality and therefore obtain an even greater understanding of the evolution of sociality.

We characterized the variability of social behavior in individual female and male prairie voles (*Microtus ochrogaster*) and its association with their mating success, reproductive success, and body condition. Prairie voles are an ideal study system to investigate sociality because both sexes can exhibit natural variation in sociality by displaying different reproductive strategies (Solomon and Jacquot, 2002). Both females and males can exhibit strategies ranging from pair bonded “residents”, with an established territory to non-territorial, unpaired “wanderers” and both are known to switch their strategy over the course of their lifetime (Getz and Hofmann, 1986; Getz et al., 1993; Solomon and Jacquot, 2002; McGuire and Getz, 2010; Shuster et al., 2019). Further, genetic monogamy and social monogamy are distinct with some female-male pairs exhibiting high levels of social and genetic monogamy, some pairs being socially monogamous but not genetically monogamous, and some individuals exhibiting no socially or genetically monogamous behavior or mating patterns (Solomon et al., 2004). Thus, prairie voles may vary in the number and strength of social associations they have with other individuals. For example, a genetically monogamous female and male likely have a very strong social association with each other whereas a wandering individual may have many weak connections with multiple opposite-sex voles. Additionally, voles may vary in the number of social connections they have with other individuals than their partner because under certain environmental conditions, such as high population density (Getz et al., 1993; Cochran and Solomon 2000; Lucia et al. 2008), voles can also form extended family groups when offspring delay dispersal from the natal nest.

We characterized the social behavior of individual prairie voles in semi-natural enclosures using an automated monitoring system. Prairie voles were all marked uniquely with passive integrated technology (PIT) tags and their movements were continuously monitored by an array of radio-frequency identification (RFID) antennas. We used patterns of spatial and temporal co-occurrence generated from this system with social network analyses to estimate patterns of social association (unweighted degree). We have previously shown that opposite-sex voles that exhibit close social associations, as generated by this automated method of data collection, are also more likely to be caught in the same trap together, exhibit overlapping home ranges, and show a strong social preference for one another in a choice experiment (Sabol et al., 2018), suggesting that these measures reflect the strength of social associations. Therefore, we classified individuals with a greater number of social network connections with opposite-sex or same-sex voles as more social, although we note that the valence of these connections (agonistic or affiliative) is not known. We characterized the sociality of all voles throughout the breeding season using social network analyses, recorded their survival through this period and used parentage analyses to quantify their mating success (number of different individuals with which they produced offspring) and reproductive success (number of offspring produced that survived to emergence from the natal nest).

We predicted that voles with a greater number of social network connections (i.e., more social) would have higher mating and reproductive success but lower body condition due to the trade-offs associated with high levels of this type of sociality. We only investigated body condition in males because any changes in body mass in females is likely directly related to pregnancy, We predicted that male voles that were more social (had more social network connections) would have lower body condition (mass) because polygyny typically involves energetic costs and risky travel associated with finding female mating partners and interacting with male competitors, which may therefore reduce survival (Blanckenhorn et al 1995; Armitage 1998).

## Methods

### Study site and study animals

All fieldwork was conducted at Miami University’s Ecology Research Center in Oxford, Ohio from May to August 2017. Voles were released into two separate 0.1 ha enclosures (33 m x 33 m). The 20-gauge sheet metal walls of the enclosures were 75 cm tall and embedded 45 cm into the ground to prevent vole movements among enclosures. Enclosure walls were topped with an electrified wire to prevent other small to medium sized mammals (e.g. raccoons and weasels) from entering the enclosures and disturbing traps. Although this system likely prevented many mammalian predators from entering the enclosures, the enclosures were open and accessible to avian predators and snakes. We had multiple animals that were confirmed to be depredated by owls and also occasionally saw large snakes within the enclosures. Prior to releasing prairie voles into the enclosures, we live-trapped within the enclosures for 3 continuous days to capture any small mammals (e.g. *Microtus pennsylvanicus*, *Peromyscus maniculatus*, or *Blarina brevicauda*) and released them outside of the enclosures.

We released laboratory-bred 7th and 8th generation prairie voles (descended from voles originally captured in Illinois) into two enclosures. The pedigree of the laboratory population was known and to avoid inbreeding, we did not place opposite-sex siblings or parents and their offspring into the same enclosure. All founding voles were sexually mature (> 31 d, Solomon 1993) but had never bred. Each enclosure was founded with a different density: Enclosure 1 was established by releasing 48 voles (24 females, 24 males) and Enclosure 2 initially contained 24 voles (12 females, 12 males). These represented densities of 480 voles/ha and 240 voles/ha respectively, which were within the range of vole densities observed in natural populations (10 to 600 voles/ha: Getz et al., 1994; Getz et al., 2001). The different starting density was employed to assess the role of density on vole social behavior but, as shown below (Fig. 1), vole density in Enclosure 1 (high density) decreased over the course of the field season. Additionally, we do not find or report an effect of density on vole social behavior in this dataset, only whether there is a difference between the enclosures in general for the whole season. The vegetation within enclosures consisted primarily of perennial grasses and forbs, which provided food and cover. Voles were not provided with supplemental food besides the cracked corn, a low-quality food, used to bait the live-traps.

**Figure 1.**
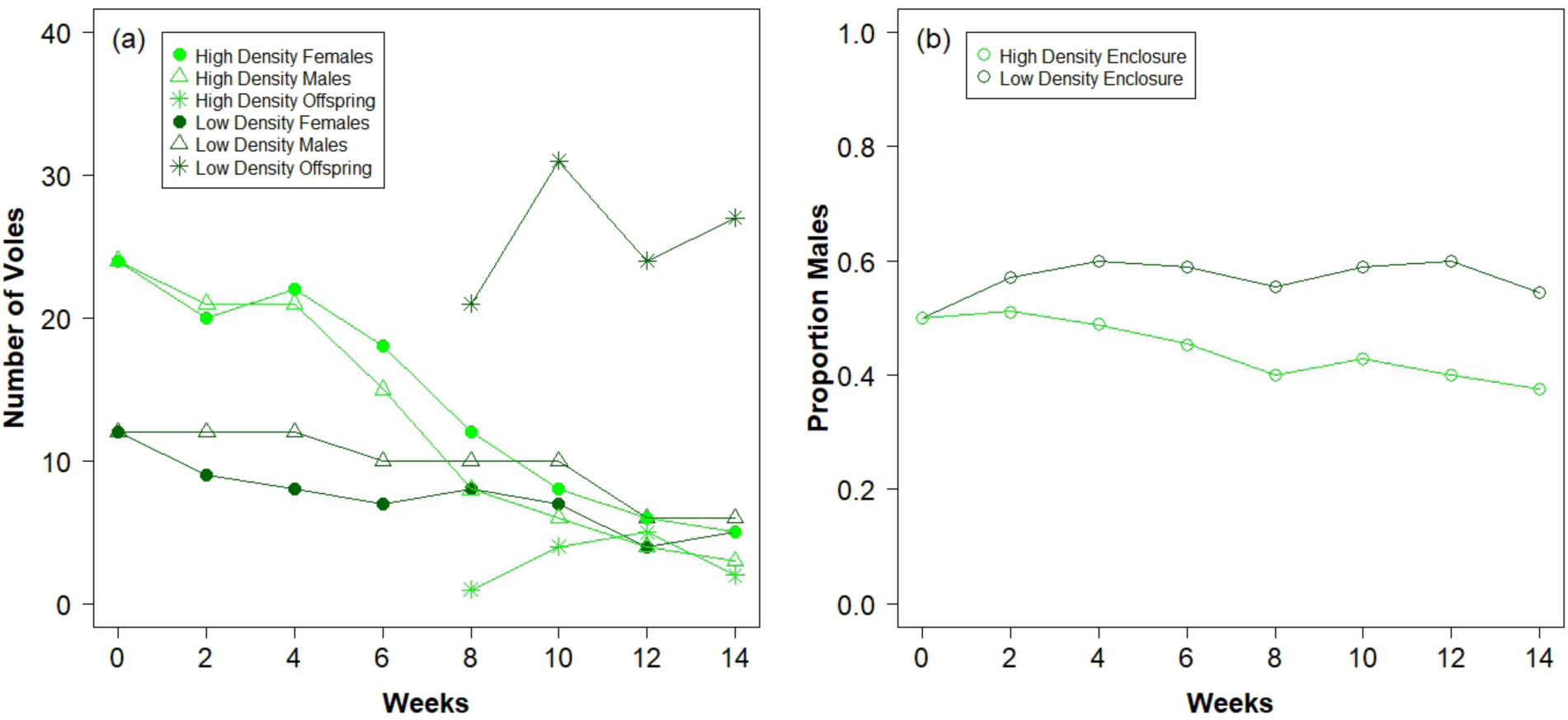
a) The number of females, males, and offspring in each of our two enclosures over time during the study based on the number of unique individuals live-trapped during each two-week period of the field season. Note that the area of enclosures is equal, so the number of voles in each enclosure can be used to compare relative density between the two, b) The sex ratio in each enclosure, calculated as the proportion of total adult voles trapped during each two-week period of the field season that were males, over time of the study.

### Recording social behavior of voles

Prior to release into the enclosures, all founding voles were implanted with a PIT tag (Biomark: Boise, Idaho) with a unique identification number. Once released, we monitored our populations through a proprietary system of 12 RFID antennas connected to a central recording station (BioMark RM310/SM303, Boise, ID) that we have used previously to create social networks of prairie voles and infer their social associations (Sabol et al., 2018). These antennas were placed within the enclosures in two different 3 x 4 arrays (Figure A1) that were rotated from array 1 to array 2 every 3 days and moved from one enclosure to the other every 6 days during the 14-week study. The antennas recorded the individual PIT tag numbers of each vole that passed within 15 cm of the antenna once a second for the entire time the animal was within this radius of the antenna. When multiple tags were within the 15 cm radius, the system alternated in recording the tag numbers so that both could be detected. This allowed us to record the natural movements and social associations of individuals in both populations, which we have previously shown to be comparable and more detailed than traditional methods of recording social associations in these populations (Sabol et al., 2018).

**Figure A1.**
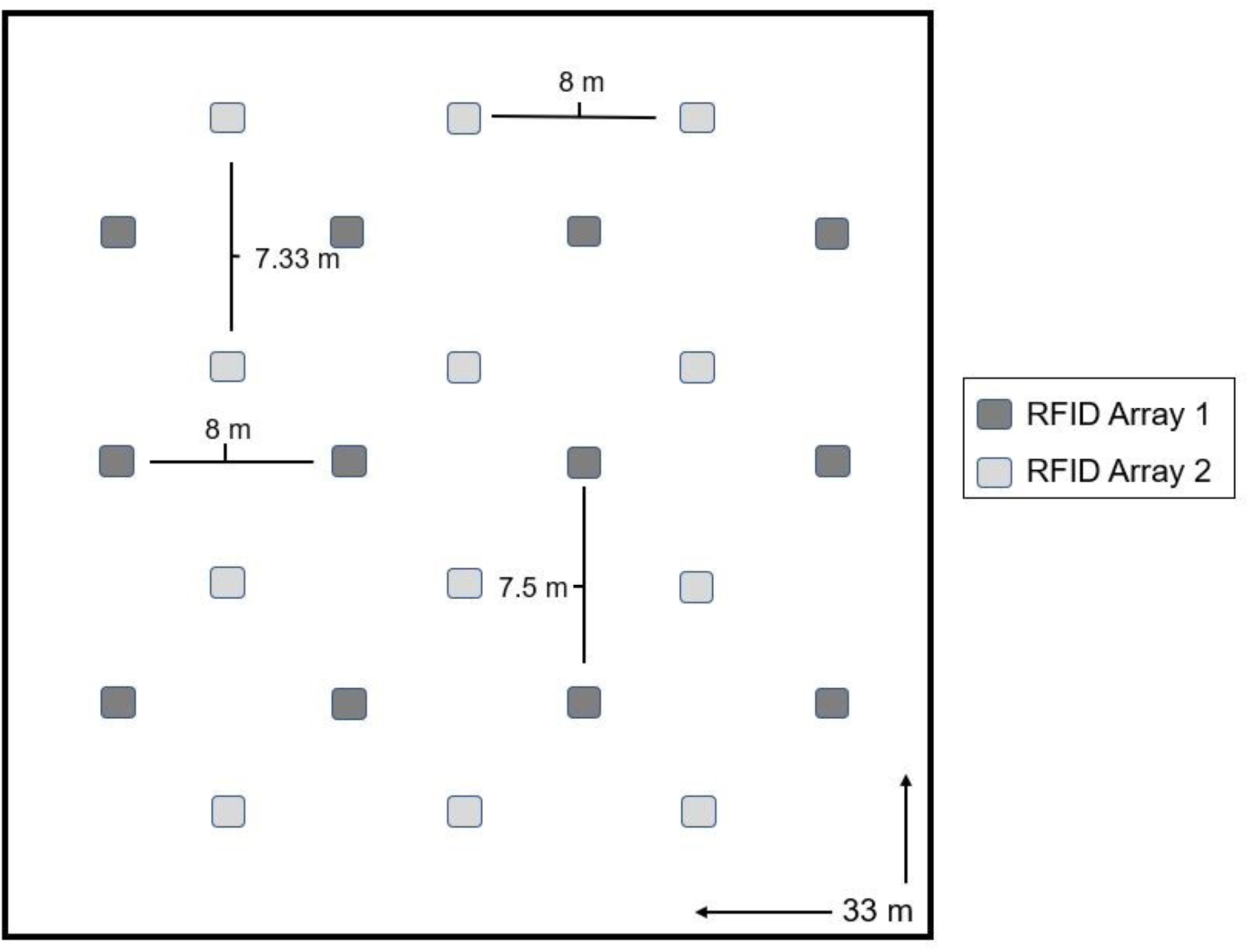
Layout of the two RFID arrays in the enclosures. The RFID system was kept at each array in each enclosure for three days in the order: array 1 enclosure 1, array 2 enclosure 1, array 1 enclosure 2, and array 2 enclosure 2 and then repeated for the duration of the field season.

### Monitoring vole reproduction

We live-trapped both enclosures by placing two Ugglan live-traps (Granhab, Hillerstorp, Sweden), baited with cracked corn, per grid stake on a 5 x 5 m trapping grid. These live-traps had a metal cover over the top to shield voles from the elements while in the traps. At the beginning of the experiment, we live-trapped nearly every day, setting traps in the evening (2230-2300 h) and checking them approximately two hours later and then leaving them set to be checked the next morning (0700 h). After the first two weeks of the experiment, we set traps approximately 3 times per week per enclosure (Monday, Wednesday, and Friday) alternating between the two enclosures so that voles in each enclosure were trapped three days over the course of two weeks. Traps were set in the evening (2230-2300 h) and checked the following morning at 0700 h. If we needed to put radiocollars on voles (see below), we also set traps from 1800-1900 h and then checked them the same evening from 2000-2100 h. Prairie vole gestation and time to weaning are each approximately 21 days (Richmond & Conaway, 1969) and, therefore, the offspring produced within the enclosures usually first emerge from the nest approximately 5-6 weeks after the adults are released (N. Solomon, B. Keane, personal observations). We therefore stopped setting traps overnight from week 6-14 of the experiment to ensure that offspring would not spend an entire night without their parents. During this time, we set traps in the evening (2230-2300 h) and checked them approximately two hours later but did not leave traps set overnight. Finally, we did not trap when there were severe thunderstorms and made up this trapping session on a different day during the week, if possible. We supplemented this regular grid trapping by placing additional traps at nest entrances after we located them using VHF telemetry and/or UV powder tracking (Lemen and Freemen, 1985).

During each capture, we identified each individual using a handheld PIT tag reader, recorded the location where the vole was live-trapped, the other individuals caught in the same trap, body mass (g, with Pesola spring scale), and assessed reproductive condition. Males were recorded as being either scrotal or non-scrotal depending on the development of the testes. Females were recorded as non-reproductive, pregnant, or lactating depending on whether developing fetuses could be felt or if nipples were pronounced. Once offspring were live trapped for the first time, we gave them a unique toe clip until they were large enough (>25 g) to be PIT tagged. Reproductive success of adult voles was estimated as the total number of offspring they produced that we were able to trap.

### Parentage analyses

Before adult voles were released into the enclosures, we collected a small piece of ear tissue and stored it in 70% ethanol in a −80° C freezer. When offspring were trapped for the first time, the tissue from the identifying toe clip was saved and temporarily stored in a −20° C freezer until samples could be moved to a −80° F freezer. We extracted DNA using DNeasy Blood and Tissue kits (Qiagen). We followed the manufacturer’s protocol except that tissue samples were incubated at 70° C, DNA was eluted in 100 µl of molecular grade water instead of 200 µl of buffer AE, and DNA samples were incubated at room temperature for 5 minutes instead of 1 minute. Once DNA was extracted, we genotyped the samples for all adults and offspring at seven microsatellite loci (Keane et al., 2007). We then ran an allele frequency analysis on the population of voles in each enclosure separately using Cervus 3.0.7 (see Mabry et al., 2011 for full details). To determine parentage, we used Cervus 3.0.7 parentage analyses with known sexes, which calculates a likelihood ratio for each potential mother and father in order to determine the most likely biological parents in the population for each offspring (Mabry et al., 2011). We were able to determine both parents (trio confidence level) with a 95% confidence level for 33/41 (80.5%) offspring, so only these 33 offspring were included in the analyses of mating and reproductive success.

### Ethical Note

All procedures involving live animals were in accordance with the guidelines provided by the American Society of Mammalogists for the use of wild mammals in research (Sikes et al., 2016) and were approved by Miami University’s Institutional Animal Care and Use Committee (protocol number 979) as this was where all work with live animals occurred.

### Statistical Analyses

All statistical analyses were done in R version 3.4.1 (R Core Team, 2017). Figure 1 was made in R while all other figures were made in ggplot2 version 2.2.1 (Wickham, 2009). All linear models and generalized linear models were run in R. For all models listed below, we assessed collinearity among the predictor variables using variance inflation factors (VIFS) in the package car, version 3.0-0 (Fox and Weisberg, 2011).

### Density and Body Mass

Population density was calculated based on the number of unique individuals caught within each two-week period (over each two-week period both enclosures were trapped with equal effort except for occasional cancellations due to weather). To investigate population density over the course of the field season, we used a linear model with density (log-transformed with base 10 to improve normality of residuals) with the fixed effects of enclosure and weeks in the study and the interaction of these terms. Sex ratio was calculated by dividing the number of adult males by the number of total adults for each two-week period. We used a binomial generalized linear model to investigate sex ratio with the fixed effects of enclosure and weeks in the study and the interaction of these terms. VIFs for all non-interaction terms were all < 3.57.

### Social Network Analyses

We measured the number of social connections (unweighted degree, hereafter degree) between same-sex or opposite-sex voles based on co-occurrence data from the RFID antennas. Individuals with a high degree would have had instances of spatial and temporal co-occurrence with many other voles whereas those with a low degree had few. We conducted all social network analyses using the R package asnipe version 1.1.4 (Farine, 2017b). In order to generate our social networks, we took the PIT tag readings from the RFID antennas and ran them through a Gaussian Mixture Model with each day labeled separately (Psorakis et al., 2012). This model goes through the raw data of the PIT tag readings and creates groups based on when tag readings at the same antenna are clumped throughout time. Therefore, there is not a uniform time period used to create these groups, they are based on how our data were distributed over time. This model uses clusters of tag readings as “centres of mass” where data are concentrated and then determines the groups based on the amount and distribution in time of tag readings in each cluster to determine where to split groups (Psorakis et al., 2012). The duration of these group events ranged from 0 seconds (so voles were both at the antenna at the same time) to 66,161 seconds with an average of 655.2 ± 3,352.8 seconds. This then creates a group by individual matrix where being in the same spatial and temporal “group” counts as an association between individuals. As we were only interested in the number of connections each individual had (not the strength of these connections), we used a binary, unweighted measurement of degree where any non-zero association was counted as a “1”. Thus, anytime we refer to the number of social connections in this paper, we calculated this using the unweighted degree. For more details about the construction of the social networks see Sabol et al. (2018).

### Reproductive Success Models

For all models including the number of mates (mating success) or the number of offspring produced that survived to emergence from the natal nest (reproductive success) as the response variable, we used Poisson generalized linear models. For each response variable we ran two models using social network data, one including all social connections in order to investigate sociality overall and one including only the opposite-sex social connections. These models all had fixed effects of the number of social connections, the interaction of this with sex, the number of social connections squared to assess non-linear effects of social connections on mating success or reproductive success (one model including all connections and another for each response variable including only opposite-sex connections), the interaction of this with sex, enclosure, and survival (calculated as the proportion of the field season the individual survived based on last detection). To test if mating success and reproductive success were related, we ran a separate model with the number of offspring produced as the response variable and fixed effects of the number of mates with which individuals produced offspring, the interaction of this with sex, and enclosure. None of the GLMs were over-dispersed as all the dispersion parameters were <1, which we tested using R package AER version 1.2.5 (Kleiber and Zeil, 2008). VIFs were all < 3.5 except interaction and squared terms, which were predictably high.

### Body Mass

To investigate body mass, we calculated the average body mass for each male vole for the entire field season (range 1-19 measurements, average 7.75 measurements). Females were not included because we were using body mass as a proxy for body quality, and female mass would be affected by both pregnancy status and body condition. We then used a general linear model for average body mass with the number of social connections (one model including all connections another including only opposite-sex connections), the number of social connections squared (one model including all connections another including only opposite-sex connections), enclosure, survival (calculated as the proportion of the field season the individual survived based on last detection), and the number of mass measurements we had for each individual. We visually assessed the distribution of the data and residuals for normality. VIFs were all < 2.2 except interaction or squared terms and survival. However, when survival was excluded from the model, VIFs for all of the other terms were < 3.5 except interactions and squared terms. Including survival did not alter the statistical significance of any of the results shown below so we left it in.

### Randomized Models

For every model that included unweighted degree (the number of social connections), we used the network permutation method in asnipe (Farine, 2017b). This method is useful because it helps control for the fact that social network data are not independent. This method also allows us to investigate our hypotheses more specifically by allowing us to test if the observed relationships are significantly different from random networks with the same structure as our social networks (see Farine et al., 2015; Spiegel et al. 2017 for other similar uses of this method). The network permutation method takes a piece of data from the group by individual matrix and swaps it for a different individual (Farine, 2013). Specifically, we ran 10,000 randomized models where each time another piece of data from the individual by group matrix was swapped. We also restricted swaps to only voles in the same enclosure that were recorded on the RFID antennas during the same day to control for voles that did not survive the entire season. Further, for the opposite-sex networks we restricted swaps to include only voles of the same sex so that we were only comparing our opposite-sex network to other opposite-sex networks, not all possible combinations. We then compared the regression coefficients from the model for each variable that includes a social network statistic to corresponding b-values from randomized networks and calculated a new *P*-value based on the number of randomized models that produced a b-value with a higher absolute value than the absolute value of the observed model. Therefore, our *P*-value shows us whether the relationship we have observed is stronger than the relationship from 10,000 randomizations of our dataset (Farine, 2013). We ran each set of randomizations three times to ensure that the *P*-values were consistently significant in each of the randomizations. We present all three *P*-values from these randomizations and, conservatively, only consider that a relationship is statistically significant if all three randomizations revealed *P-*values <0.05.

## Results

### Enclosure density & adult sex ratio

The number of adult voles in each enclosure declined over the course of the field season due to mortality (effect of weeks in the study, b = −0.14, SE = 0.012, t_12_ = −11.45, *P* < 0.0001, Fig. 1a, note estimates on log_10_ scale) and the significant interaction between weeks in the study and enclosure indicated that vole density decreased more strongly in the high density enclosure than in the low density enclosure (enclosure x weeks: b = 0.083, SE = 0.017, t_12_ = 4.79, *P* = 0.00045). For example, the starting density of voles in the high-density enclosure was 480 voles/ha (week 0 in Fig. 1a) but was reduced to 200 voles/ha in the middle of the experiment (week 8 in Fig. 1a) and to 60 voles/ha at the end of the experiment (week 14 in Fig. 1a). The starting density in the low-density enclosure was 240 voles/ha (week 0 in Fig. 1a) but was reduced to 180 voles/ha in the middle of the experiment (week 8 in Fig. 1a) and 100 voles/ha at the end of the experiment (week 14 in Fig. 1a). In total, only 12.5% of voles in the high-density enclosure were still alive at the end of the experiment whereas 41.7% of the voles in the low-density enclosure were still alive at the end of the experiment.

Unlike density, adult sex ratio did not differ significantly during the course of the experiment (effect of time, b = −0.04, SE = 0.16, z = −0.26, df = 12, *P* = 0.80, Fig. 1b) or between the two enclosures (effect of enclosure, b = 0.15, SE = 1.84, z = 0.082, df = 12, *P* = 0.94, Fig. 1b).

### Effects of sociality on mating success

Overall, both female and male voles that had an intermediate (i.e., the middle of the range of observed values) number of social connections produced offspring with a greater number of different mates (i.e., had higher mating success). In the model considering all social interactions with same- and opposite-sex individuals, voles that had an intermediate number of social connections (degree) with all possible individuals had higher mating success (effect of social connections^2^: b = −0.012, z = −0.88, *P*-values from randomized networks = 0.012, 0.0093, <0.0001, Table 1, Fig. 2a) but this relationship was slightly different between the sexes (sex x social connections^2^: b = 0.014, z = 0.94, *P* = 0.010, 0.0018, <0.0001, Table 1, Fig. 2a). In both females and males, those with an intermediate number of social connections had the highest mating success, therefore the interaction with sex and the number of social connections^2^ on mating success seemed to be largely due to males having slightly more overall social connections than females while female mating success peaked at a lower number of social connections (Fig. 2a). There is also a qualitative difference in the shape of the curve, with female mating success peaking at a lower number of social connections but then dropping off more steeply, while male mating success peaked at a higher number of social connections but declined more gradually (Fig. 2a).

**Figure 2.**
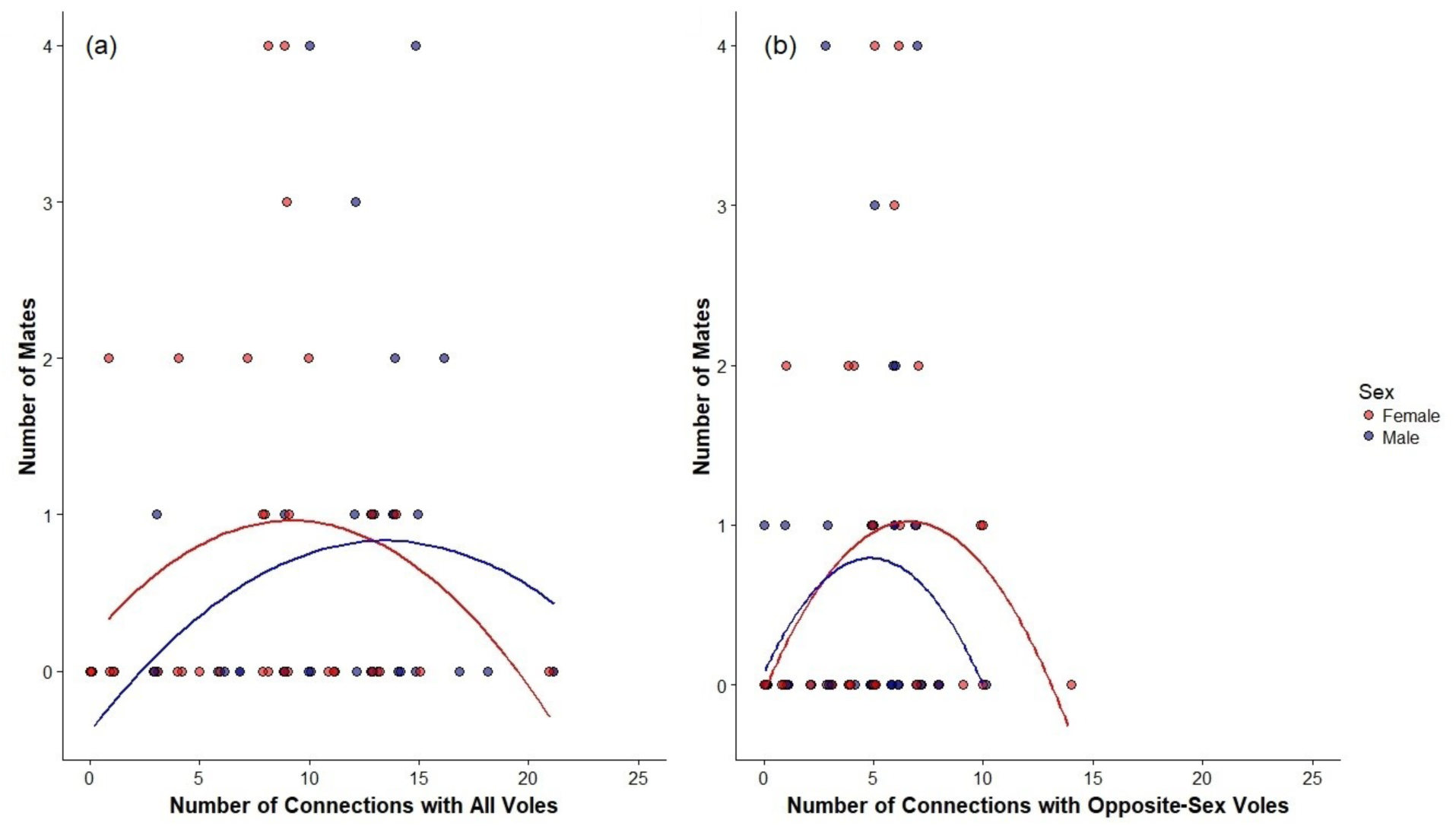
Both females and males with an intermediate number of social connections with a) all voles in their enclosure (both same- or opposite-sex individuals) or b) only opposite-sex individuals in their enclosure had higher mating success (defined as the number of different individuals with which they produced offspring). Points are jittered with males shown in blue and females in red. Full statistical results shown in Table 1.

**Table 1.** Effects of the number of social connections on vole mating success (number of different individuals a vole produced offspring with).

| Social Network | Variable | Estimate | SE | z-value | P-value from GLM | P-values from Randomization Tests |
| --- | --- | --- | --- | --- | --- | --- |
| All individuals | Intercept | -2.66 | 0.65 | -4.06 | <0.0001 | NA |
|  | Sex (M) | 0.68 | 0.92 | 0.73 | 0.46 | NA |
|  | Enclosure (LD) | 1.89 | 0.41 | 4.55 | <0.0001 | NA |
|  | Survival | 2.07 | 0.89 | 2.33 | 0.020 | NA |
|  | Degree | 0.083 | 0.21 | 0.40 | 0.69 | 0.12, 0.0098, 0.14 |
|  | Degree<sup>2</sup> | -0.012 | 0.013 | -0.88 | 0.38 | 0.012, 0.0093, <0.0001 |
|  | Sex (M) x Degree | -0.21 | 0.23 | -0.90 | 0.37 | 0.021, 0.0017, <0.0001 |
|  | Sex (M) x Degree<sup>2</sup> | 0.014 | 0.015 | 0.94 | 0.35 | 0.01, 0.0018, <0.0001 |
| Opposite-sex individuals | Intercept | -2.81 | 0.70 | -4.08 | <0.0001 | NA |
|  | Sex (M) | 0.78 | 0.91 | 0.86 | 0.39 | NA |
|  | Enclosure (LD) | 2.03 | 0.43 | 4.72 | <0.0001 | NA |
|  | Survival | 1.75 | 0.84 | 2.09 | 0.037 | NA |
|  | Degree | 0.19 | 0.31 | 0.63 | 0.53 | 0.18, 0.19, <0.0001 |
|  | Degree<sup>2</sup> | -0.028 | 0.028 | -1.01 | 0.31 | 0.0011, 0.0035, 0.001 |
|  | Sex (M) x Degree | -0.53 | 0.38 | -1.42 | 0.15 | <0.0001, <0.0001, 0.0046 |
|  | Sex (M) x Degree<sup>2</sup> | 0.057 | 0.042 | 1.36 | 0.17 | <0.0001, <0.0001, 0.0068 |

Degree refers to the number of social connections for each individual either with all voles or opposite-sex voles. Note that relationships involving social network data have three *P*-values because those regression coefficients were compared to those from randomized networks three times to determine if they were consistently significant (see methods for details). Survival refers to the proportion of days the vole was in the enclosure based on when it was last recorded. “LD” is low-density enclosure.

The same relationship was true when only opposite-sex connections were considered. Mating success was highest for female and male voles with an intermediate number of opposite-sex social connections (social connections^2^: b = −0.028, z = −1.01, *P* = 0.0011, 0.0035, 0.001, Fig. 2b), although the magnitude of this effect slightly differed between the sexes (sex x social connections^2^: b = 0.057, z = 1.36, *P* = <0.0001, <0.0001, 0.0068, Table 1, Fig. 2b). This latter difference between the sexes seems to be driven by the difference in the number of social connections between the sexes with females tending to have slightly more social connections than males.

Overall, both female and male voles in the low-density enclosure had higher mating success than individuals in the higher density enclosure (from model for all social connections: b = 1.89, z = 4.55, *P* < 0.0001; from model for all opposite-sex social connections: b = 2.03, z = 4.72, *P* < 0.0001, Table 1). Individuals that survived in the enclosures for longer had higher mating success (all social connections: b = 2.07, z = 2.33, *P* = 0.020; opposite-sex social connections: b = 1.75, z = 2.09, *P* = 0.037, Table 1).

### Effects of sociality on reproductive success

Both female and male voles with an intermediate number of social connections with all voles produced more offspring that survived to emergence from the natal nest (b = −0.0085, z = −0.77, *P* = 0.002, 0.035, 0.015, Table 2, Fig. 3a). This result did not consistently differ by sex (sex x social connections^2^: b = 0.0082, z = 0.63, *P* = 0.021, 0.11, 0.022, Table 2, Fig. 3a). However, when only considering opposite-sex social connections (Fig. 3b), these relationships were not significant. The number of offspring that voles produced was not related to the number of opposite-sex social connections for males or females (effect of opposite-sex social connections: b = 0.15, z = 0.52, *P* = 0.20, 0.53, 0.54, Table 2, Fig. 3b) and voles with an intermediate number of opposite-sex social connections did not produce significantly more offspring (effect of opposite-sex social connections^2^: b = −0.022, z = −0.93, *P* = 0.14, 0.099, 0.12, Table 2, Fig. 3b). This relationship did not consistently vary with sex, as the difference between the sexes for opposite-sex connections^2^ where the inverted u-shaped relationship was slightly lessened in males and was not significant in all three sets of randomizations (sex x opposite-sex connections^2^: b = 0.038, z-value = 0.92, *P* = 0.063, 0.010, 0.012, Table 2, Fig 3b) and the difference between opposite sex connections between males and females where males tended to have fewer connections than females was also not significant in all three sets of randomizations (sex x number of opposite-sex connections: b = −0.39, z = −1.04, *P* = 0.047, 0.024, 0.11). Male and female voles that survived for longer produced more offspring (all social connections: b = 2.49, z = 3.03, P = 0.0025; opposite-sex social connections: b = 2.19, z = 2.83, P = 0.0046, Table 2).

**Figure 3.**
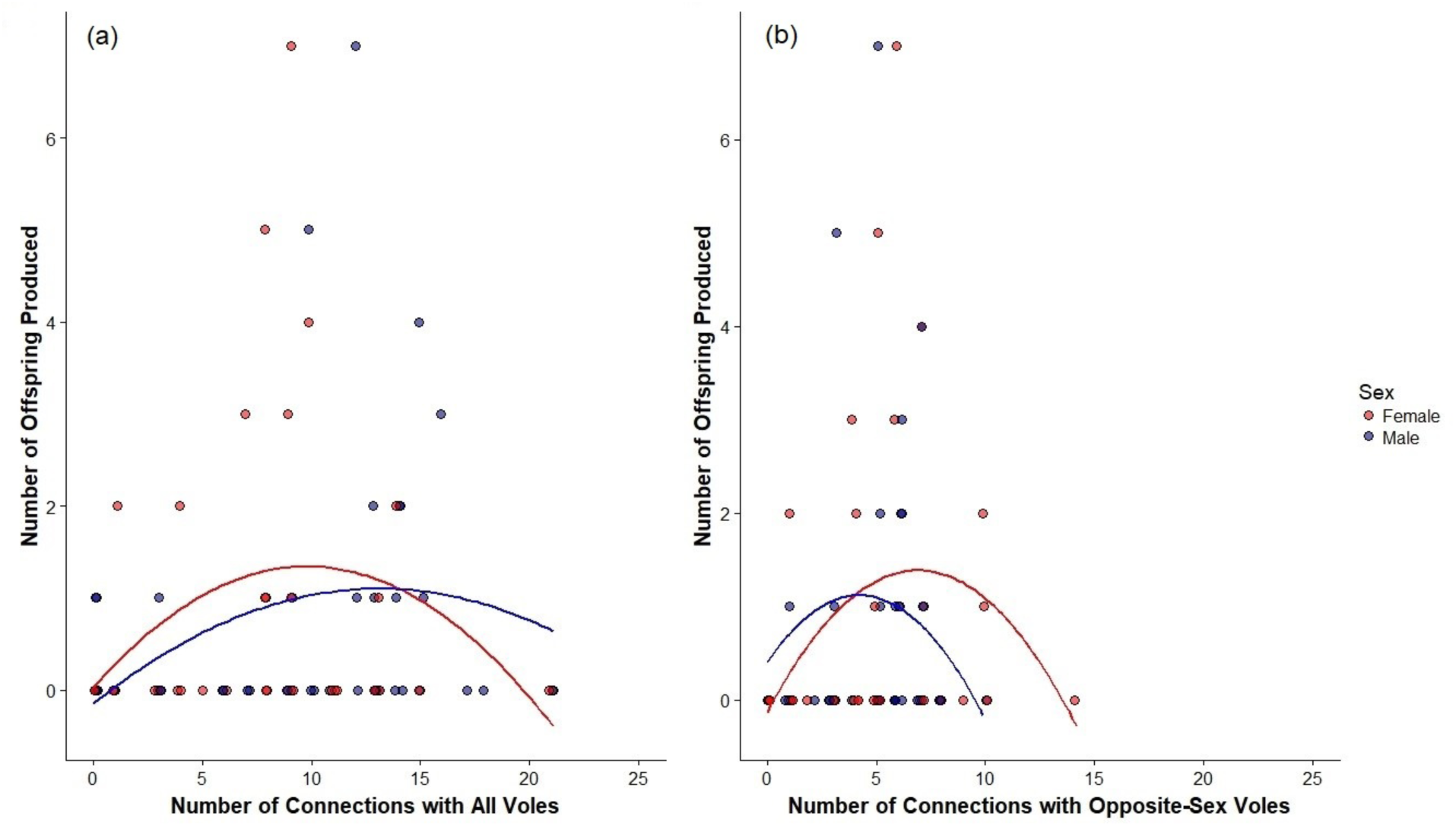
a) Both females and males with an intermediate number of social connections with all voles in their enclosure (both same- and opposite-sex individuals) produced more offspring that survived to emergence from the natal nest. b) Female and male voles with an intermediate number of social connections with only opposite-sex individuals in their enclosure also tended to produce more offspring. Points are jittered with red points being females and blue points being males. Full statistical results shown in Table 2.

**Table 2.** Effects of the number of social connections on vole reproductive success (number of offspring produced that survived until emergence from the natal nest).

| Social Network | Variable | Estimate | SE | z-value | P-value from GLM | P-values from Randomization Tests |
| --- | --- | --- | --- | --- | --- | --- |
| All individuals | Intercept | -2.90 | 0.66 | -4.14 | <0.0001 | NA |
|  | Sex (M) | 0.61 | 0.93 | 0.65 | 0.52 | NA |
|  | Enclosure (LD) | 2.10 | 0.40 | 5.24 | <0.0001 | NA |
|  | Survival | 2.49 | 0.82 | 3.03 | 0.0025 | NA |
|  | Degree | 0.053 | 0.19 | 0.28 | 0.78 | 0.33, 0.25, 0.19 |
|  | Degree<sup>2</sup> | -0.0085 | 0.011 | -0.77 | 0.44 | 0.002, 0.035, 0.015 |
|  | Sex (M) x Degree | -0.14 | 0.21 | -0.64 | 0.52 | 0.12, 0.18, 0.12 |
|  | Sex (M) x Degree<sup>2</sup> | 0.0082 | 0.013 | 0.63 | 0.53 | 0.021, 0.11, 0.022 |
| Opposite-sex individuals | Intercept | -2.97 | 0.67 | -4.41 | <0.0001 | NA |
|  | Sex (M) | 0.58 | 0.91 | 0.64 | 0.53 | NA |
|  | Enclosure (LD) | 2.21 | 0.41 | 5.40 | <0.0001 | NA |
|  | Survival | 2.19 | 0.77 | 2.83 | 0.0046 | NA |
|  | Degree | 0.15 | 0.28 | 0.52 | 0.60 | 0.20, 0.53, 0.54 |
|  | Degree<sup>2</sup> | -0.022 | 0.024 | -0.93 | 0.35 | 0.14, 0.099, 0.12 |
|  | Sex (M) x Degree | -0.39 | 0.38 | -1.04 | 0.30 | 0.047, 0.024, 0.11 |
|  | Sex (M) x Degree<sup>2</sup> | 0.038 | 0.041 | 0.92 | 0.36 | 0.063, 0.010, 0.02 |

Degree refers to the number of social connections for each individual either with all voles or opposite-sex voles. Note that relationships involving social network data have three *P*-values because those regression coefficients were compared to those from randomized networks three times to determine if they were consistently significant (see methods for details). Survival refers to the proportion of days the vole was in the enclosure based on when it was last recorded. “LD” is low-density enclosure.

### Effects of mating tactics on reproductive success

Overall, there was a fitness advantage to being polygynous (mating with a greater number of individuals) in both females and males. Individuals with more mates produced more offspring that survived to emergence from the natal nest (b = 0.64, z = 4.88, *P* <0.0001, Table 3, Fig. 4) and this relationship did not differ by sex as the interaction between number of mates and sex was not significant (b = −0.093, z = −0.56, *P* = 0.58, Table 3). Individuals in the lower density enclosure had higher reproductive success than individuals in the higher density enclosure (b = 1.53, z = 3.37, *P* = 0.00076, Table 3).

**Figure 4.**
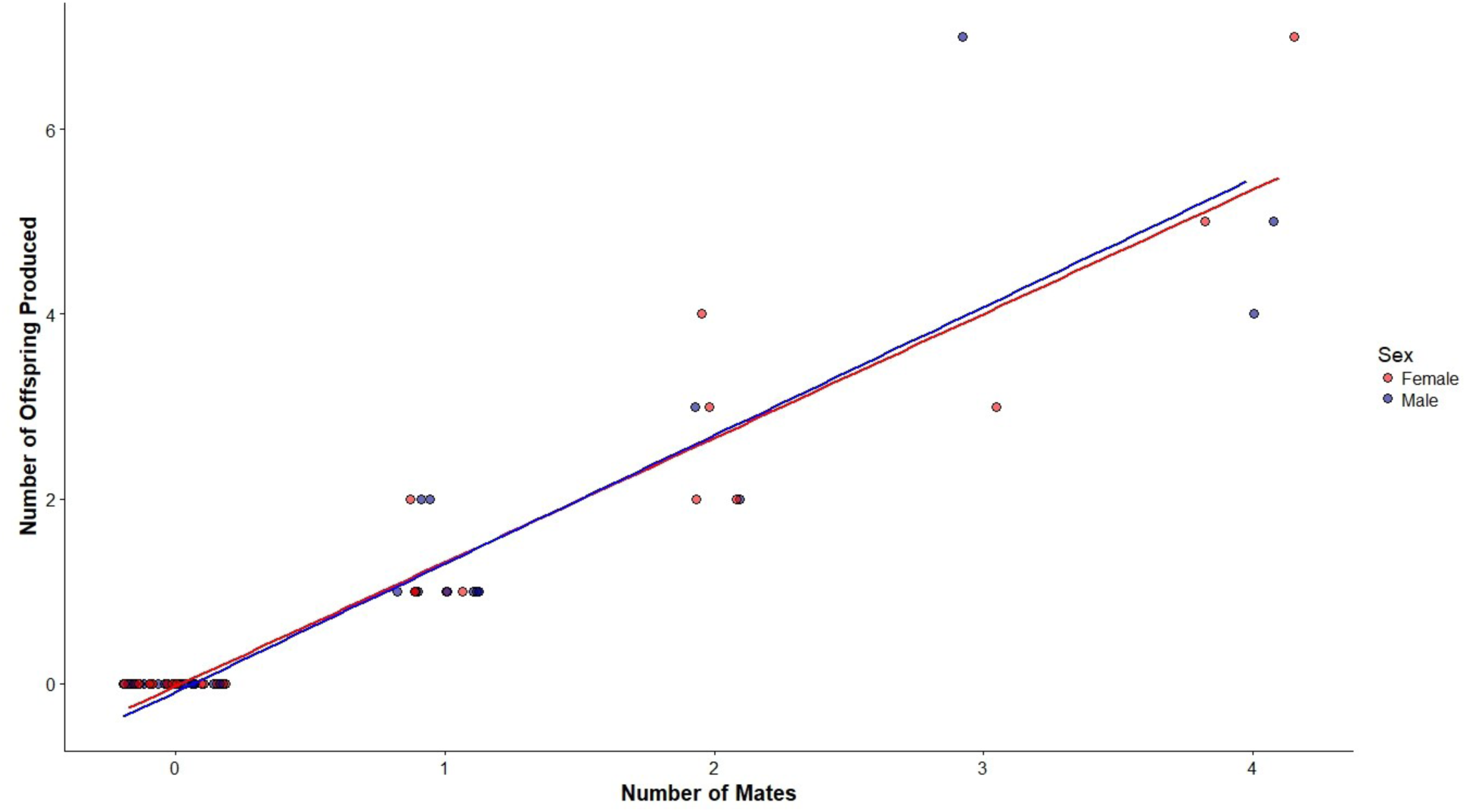
Both female and male prairie voles that had higher mating success (produced offspring with a greater number of different mates) produced a great number of offspring that survived to emergence from the natal nest. Points for females and males are jittered. Full results shown in Table 3.

**Table 3.** Overall effects of vole mating success (number of different individuals with which a vole produced offspring) on reproductive success (number of offspring produced that survived until emergence from the natal nest).

| Variable | Estimate | SE | z-value | P-value from GLM |
| --- | --- | --- | --- | --- |
| **Intercept** | **-2.13** | **0.44** | **-4.83** | **<0.0001** |
| **Number of Mates** | **0.64** | **0.13** | **4.88** | **<0.0001** |
| Sex (M) | 0.26 | 0.49 | 0.53 | 0.60 |
| **Enclosure (LD)** | **1.53** | **0.45** | **3.37** | **0.00076** |
| # of Mates x Sex (M) | -0.093 | 0.17 | -0.56 | 0.58 |

Since this model did not include social network data we did not perform the randomizations to generate three *P*-values, we present the *P*-values from the GLM. “LD” is low-density enclosure.

### Effects of sociality on body mass

Males with an intermediate number of connections had a higher average body mass (effect of social connections^2^: b = −0.066, t = −2.36, *P* = <0.0001, <0.0001, <0.0001, Table 4, Fig. 5a). This relationship was also significant when only opposite-sex connections were considered (effect of opposite-sex social connections^2^: b = −0.20, t = 1.54, *P* = 0.0024, 0.0031, 0.0005, Table 4, Fig. 5b). Average body mass was not significantly different between the two enclosures (all social connections: b = −1.03, t = −0.40, *P* = 0.69; opposite-sex social connections: b = −0.16, t = −0.058, *P* = 0.95, Table 4). Survival did not predict average body mass in either model (all social connections: b = 13.37, t = 1.52, P = 0.14; opposite-sex social connections: b = 10.80, t = 1.22, *P* = 0.23, Table 4). Although the number of measurements we obtained to calculate each individual’s average body mass varied (Fig. 5), it did not affect our measure of average body mass in either model (all social connections: b = −0.12, t = −0.20, *P* = 0.84; opposite-sex social connections: b = −0.20, t = −0.31, *P* =0.76, Table 4).

**Figure 5.**
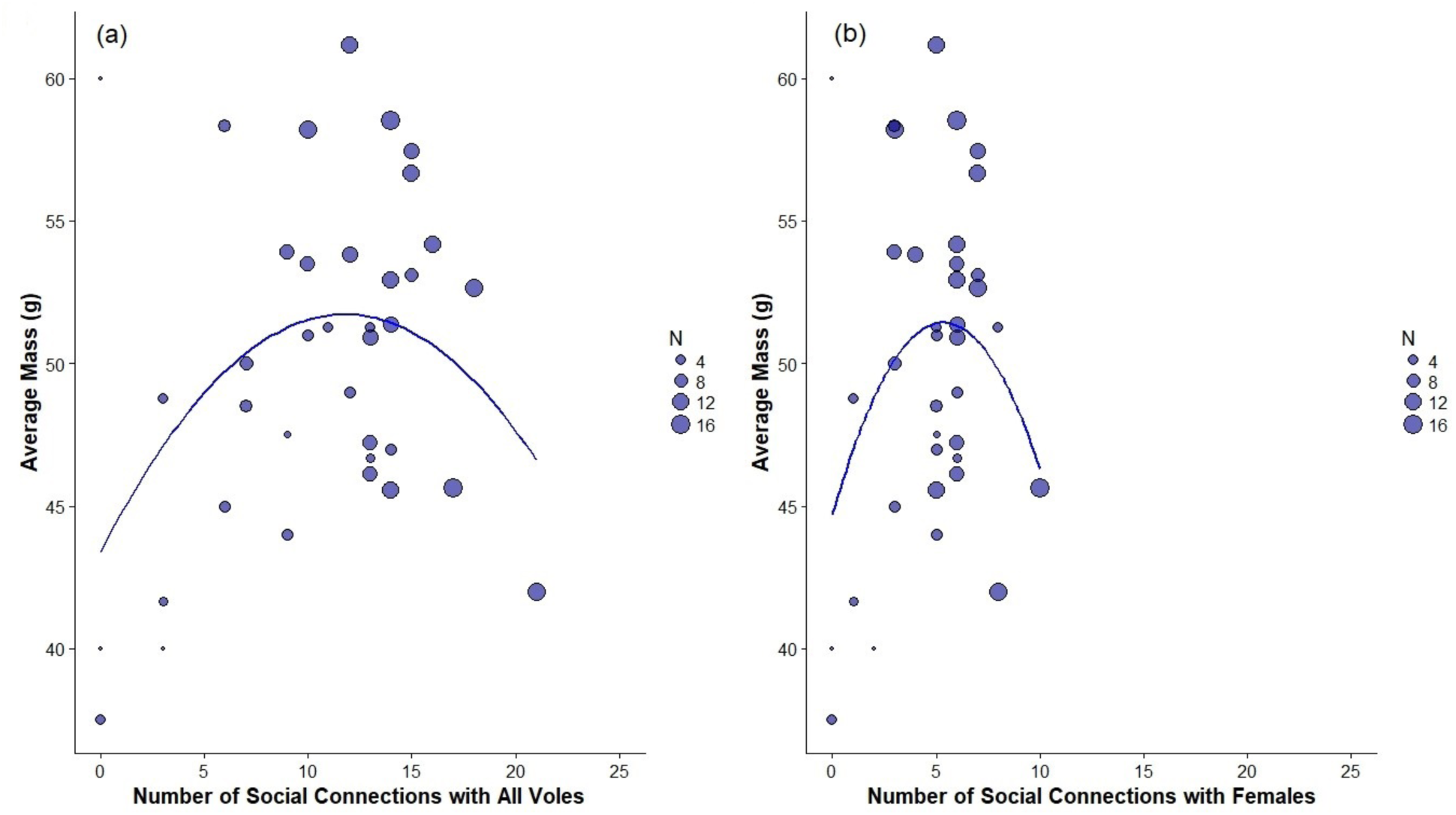
Male voles with an intermediate number of social connections a) with both female and male voles in their enclosure or b) with just female voles, were significantly heavier over the course of this study. Body mass for males was averaged for the entire duration of this study. The number of times we measured body mass (“N”) varied among males so the size of each point is scaled based on the number of recorded mass measurements we have for each individual. Full results shown in Table 4.

**Table 4.** Effects of the number of social connections on average body mass for males.

| Social Network | Variable | Estimate | SE | t-value | P-value from LM | P-values from Randomization Tests |
| --- | --- | --- | --- | --- | --- | --- |
| **All individuals** | **Intercept** | **43.42** | **2.70** | **16.10** | **<0.0001** | NA |
|  | Enclosure (LD) | -1.03 | 2.57 | -0.40 | 0.69 | NA |
|  | Survival | 13.37 | 8.78 | 1.52 | 0.14 | NA |
|  | # of measures | -0.12 | 0.60 | -0.20 | 0.84 | NA |
|  | **Degree** | **0.95** | **0.56** | **1.70** | **0.099** | **<0.0001, 0.0001, <0.0001** |
|  | **Degree<sup>2</sup>** | **-0.066** | **0.028** | **-2.36** | **0.025** | **<0.0001, <0.0001, <0.0001** |
| **Opposite-sex individuals** | **Intercept** | **44.31** | **2.81** | **15.78** | **<0.0001** | NA |
|  | Enclosure (LD) | -0.16 | 2.80 | -0.058 | 0.95 | NA |
|  | Survival | 10.80 | 8.89 | 1.22 | 0.23 | NA |
|  | # of measures | -0.20 | 0.63 | -0.31 | 0.76 | NA |
|  | **Degree** | **1.54** | **1.22** | **1.26** | **0.22** | **0.0035, 0.0036, 0.0332** |
|  | **Degree<sup>2</sup>** | **-0.20** | **0.13** | **-1.54** | **0.13** | **0.0024, 0.0031, 0.0005** |

Degree refers to the number of social connections for each individual either with all voles or opposite-sex voles. Relationships involving social network data have three *P*-values because those regression coefficients were compared to those from randomized networks three times to determine if they were consistently significant (see methods). “# of measures” refers to number of times we measured body mass, which were used to generate average mass for each individual. Survival refers to proportion of days the vole was in the enclosure. “LD” is low-density enclosure.

## Discussion

As expected, both female and male prairie voles varied in the frequency of social interactions, as reflected in our social network analyses that quantified social network degree using the frequency of temporal and spatial co-occurrence generated by our RFID system. Female and male voles with an intermediate number of social interactions had the greatest mating success and produced the greatest number of offspring, though the latter was only the case when we included all interactions with other voles, and not when only opposite-sex connections were considered.

We tested our predictions about the fitness benefits of sociality using two different sets of social network data, one including social connections with all voles and one only including connections with opposite-sex individuals. This allowed us to investigate the potential costs and benefits of sociality overall (e.g., social interactions of a female vole with other females and males) as well as specifically social interactions between opposite-sex individuals, which may be more directly relevant to mating and reproductive success. We found that voles with an intermediate number of social connections with all voles (i.e., more social) had significantly higher mating and reproductive success whereas voles with an intermediate number of opposite-sex connections had significantly higher mating success but not higher reproductive success. The process of running the randomizations was somewhat different for the two sets of networks (all social connections or just opposite-sex connections), which could contribute to the observed differences in the results between the two sets of networks. We limited permutations for the opposite-sex network to within the same sex (where female social network data were being swapped for another female and male data were being swapped for another male) so these swaps could only be done between approximately half as many individuals each time. By keeping the sex of each individual in the association consistent, the structure of the randomizations was more like the data it was being compared to instead of comparing all possible connections (Farine, 2017a). However, since the permutations were done on the raw data and then we pulled only the opposite-sex connections from these networks, it is possible that some of the 10,000 permutations affected the same-sex connections in the raw data (therefore changing a social interaction that is not included in data used for the linear model) and therefore may not have changed the estimate for the relationship between opposite-sex connections and the response variable every time, whereas in the models with all connections included, a relevant social interaction would have changed every time, which would then change the model estimate for the relationship between social connections and the response variable some amount every time. Since we ran such a large number of these permutations, this may not have affected the overall result of the tests, but it is a limitation of the method.

Using our measure of sociality, our results suggest the possibility of stabilizing selection on sociality because voles that co-occurred spatially and temporally with very few or very many conspecifics had the lowest mating success and, at least when considering all social connections, had the lowest reproductive success. Both McGuire et al. (2002) and Solomon and Keane (2018) showed that large social groups do not increase female reproductive success. McGuire et al. (2002) also showed that female prairie voles that lived in large groups had fewer offspring survive to 12 or 30 days of age. Similarly, Solomon and Keane (2018) showed that females did not benefit from living in large social groups in two other natural populations. These studies are consistent with our results that a very large number of social connections (which should occur in large social groups) does not increase reproductive success of breeding females, and our results show that this is also true for males.

We only recorded the association between one measure of sociality and mating and reproductive success in one year and the effects of the number of social connections on fitness could be altered when environmental conditions change. For example, the Female Dispersion Hypothesis would predict that if our measure of sociality reflects the socially monogamous behavior of male prairie voles, our measure of sociality should be positively correlated with male mating success when females are spatially clumped as males that have more social connections with clumped females should have higher mating success (Shuster and Wade, 2003; Dobson et al., 2010; Lukas and Clutton-Brock, 2013). In natural populations of prairie voles, density is quite variable across years (Getz et al., 1993, 2001) and some previous observational studies of prairie voles in field settings suggested that socially monogamous behavior is more common at low densities (McGuire et al.,1990; Solomon et al., 2009; but see Getz and McGuire,1993). There is also some evidence that resource distribution may impact the mating strategy of prairie voles and this effect may be mediated through its influence on density (Streatfeild et al., 2011). This suggests the possibility that selection on the social behavior of prairie voles varies among years due to changes in population or female density but additional multi-year studies measuring a broader array of social behaviors in free-living voles are needed to test this prediction.

One possible explanation for an intermediate level of sociality being associated with the highest mating success and potentially highest reproductive success is that this reflects a tradeoff between devoting time to social interactions (although we do not know the type of social interaction occurring) with other voles and time to other behaviors like foraging and parental care. Although high levels of sociality can have beneficial effects on individual fitness, it may also carry costs for an individual’s health or physical condition (Nunn et al., 2015). Indeed, we found that male voles with the most social connections had the lowest body mass, suggesting that there may be a reduction in body condition associated with a very high level of sociality. This could reflect the energetic costs associated with having many social connections or living in a large group (e.g., Lutermann et al., 2013), or these could be agonistic interactions with males on neighboring territories, resulting in males investing more time in territory defense than males with fewer neighbors. Why males with very few social connections were also lighter in body mass is not clear but these males may have been of lower phenotypic quality given that they had few social connections, low body mass, and low mating and reproductive success. Alternatively, having fewer social connections could result in a loss of body mass if these males had no assistance in territory defense and thus, expended more energy than males with more social connections (e.g., having a female social partner). Females likely face many of these same tradeoffs, but as we did not test quality in females (due to changes in mass being linked to pregnancy) and so further study is needed to investigate this relationship in females.

Our results suggest that it is not advantageous for voles to have social connections with too many opposite-sex conspecifics. One possible explanation is that individuals with an intermediate number of social connections may better balance the trade-off between the number and quality or strength of social relationships. For example, individuals with the highest social network degree may just have many weak social or agonistic connections, which may not result in more matings or increased reproductive success. Individuals with an intermediate social network degree may have more affiliative social connections that are strong enough to result in matings than voles at either extreme. This also is reflected in the fact that voles have many more social connections than actual mating partners (Fig. 2), where the range of the number of mates varies from 0 to 4 while the range of the number of social connections with opposite-sex individuals is from 0 to 14. This is supported by studies of the association between the strength of social connections and fitness in cercopithecine primates (baboons) where females with strong social bonds with other females in their group have higher offspring survival (Silk et al., 2003, 2009) or longevity (Silk et al., 2010). Similar relationships between the strength of social bonds and fitness have also been found in male primates; male Assamese macaques (*Macaca assamensis*) with strong social bonds to other males (including unrelated males) sired more offspring than those with fewer strong bonds with other males (Schülke, et al., 2010). As the number of social connections increase, the strength of association of each of these social connections may decline (Whitehead 2008), thus prairie voles may be constrained by the number of social connections in which they can invest enough time to result in successful mating or rearing of offspring given that prairie voles exhibit biparental care. Individuals that can best balance this trade-off between the number and strength of social connections may have the highest mating and reproductive success.

It is of course likely that the fitness benefits of the quantity versus quality of social connections may vary according to whether the modal social structure of the species is group-living (such as primate species mentioned above where strong social bonds increase fitness) or its mating system. For example, Ryder et al. (2009) found a positive association between the number of social connections (social network degree) and number of offspring sired in male manakins. As this is a lekking species, coordinated male displays may make male-male connections a more important factor for mating success than in prairie voles. Additionally, the short-term coalitions at leks may make the strength of the relationship less important than in species like prairie voles. Studies like these are rare and so future studies across a broader array of species with different mating systems will be needed to fully characterize the relationship between the number and strength of social connections and measures of fitness. Doing so will help provide insight into how individuals within a species balance the fitness costs and benefits of social behavior, thereby providing a complementary approach to comparative studies regarding the evolution of social behavior.

## Acknowledgements

We thank the Ecology Research Center (ERC) at Miami University for permitting us to conduct this study and Jeremy Fruth, field manager at the ERC, for coordinating the project. We also thank James Lichter, Sage Sparks, Caleb Collins, Tony Leon, Meghan Charles, and Staci Burrow for assisting on the project. We also thank the American Society of Mammalogists (Grant-in-Aid of Research to A. Sabol) and University of Michigan (B. Dantzer) for funding.

## Data Statement

All raw data and R code are available under CC-BY from FigShare at the link https://figshare.com/projects/How_does_individual_variation_in_sociality_influence_fitness_in_prairie_voles_/71969).

## Literature Cited

Alexander, R. D. (1974). The evolution of social behaviour. Annual Review of Ecology, Evolution, and Systematics, 5, 325–383.

Armitage, K. B. (1998). Reproductive strategies of yellow-bellied marmots: Energy conservation and differences between the sexes. Journal of Mammalogy, 79, 385–393.

Blanckenhorn, W. U., Preziosi, R. F. & Fairbairn, D. J.(1995). Time and energy constraints and the evolution of sexual size dimorphism — to eat or to mate? Evolutionary Ecology, 9, 369–381. doi:10.1007/BF01237760

Clutton-Brock, T. H. (1989). Review lecture: mammalian mating systems. Proceedings of the Royal Society B, 236, 339–372.

Clutton-Brock, T.H., Brotherton, P.N.M., Smith, R., McIllrath, G.M., Kansky, R., Gaynor, D., O’rian, M.J., Skinner, J.D. (1998). Infanticide and expulsion of females in a cooperative mammal. Proceedings of the Royal Society B, 265, 2291–2295.

Cochran, G. R., & Solomon, N. G. (2000). Effects of food supplementation on the social organization of prairie voles (*Microtus ochrogaster*). Journal of Mammalogy, 81, 746–757.

Côté, I.M., & Poulinb R. (1995). Parasitism and group size in social animals: a meta-analysis. Behavioral Ecology, 6, 159–165.

Creel, S., Dantzer, B., Goymann, W. & Rubenstein, D.R. (2013). The ecology of stress: effects of the social environment. Functional Ecology, 27, 66–80.

Dantzer, B., Goncalves, I.B., Spence-Jones, H.C., Bennett, N.C., Heistermann, M., Ganswindt, A., Dubuc, C., Gaynor, D., Manser, M.B., Clutton-Brock, T.H. (2017). The influence of stress hormones and aggression on cooperative behaviour in subordinate meerkats. Proceedings of the Royal Society B, 284, 20171248.

Dobson, F. S., Way, B. M., & Baudoin C. (2010). Spatial dynamics and the evolution of social monogamy in mammals. Behavioral Ecology, 21, 747–752. doi:10.1093/beheco/arq048

Donaldson, Z. R., & Young, L. J. (2008). Oxytocin, vasopressin, and the neurogenetics of sociality. Science, 322, 900–904. doi:10.1126/science.1158668

Eisenberg, J.F., Muckenhirn, N.A., Rudran, R. (1972). The relation between ecology and social structure in primates. Science, 176, 863–874.

Emlen, S.T. (1984). Cooperative breeding in birds and mammals. Behavioural Ecology: An Evolutionary Approach. J.R. Krebs and N.B. Davies (Eds.), pp. 305–339, Blackwell, Oxford.

Emlen, S.T. (1994). Benefits, constraints and the evolution of the family. Trends in Ecology and Evolution, 9, 282–285.

Ewald, P. W. (1994) Evolution of Infectious Disease. Oxford: Oxford University Press.

Farine, D. R. (2013). Animal social network inference and permutations for ecologists in R using *asnipe*. Methods in Ecology and Evolution, 4, 1187–1194. doi:10.1111/2041-210X.12121

Farine, D. R., Firth, J. A., Aplin, L. M., Crates, R. A., Culina, A., Garroway, C.J., Hinde, C. A., Kidd, L. R., Milligan, N. D., Psorakis, I., Radersma, R., Verhelst, B., Voelkl, B., & Sheldon, B. C. (2015). The role of social and ecological processes in structuring animal populations: a case study from automated tracking of wild birds. Royal Society Open Science, 2. doi:10.1098/rsos.150057

Farine, D. R. (2017a). A guide to null models for animal social network analysis. Methods in Ecology and Evolution, 8, 1309–1320. doi:10.1111/2041-210X.12772

Farine, D. R. (2017b). *asnipe: Animal Social Network Inference and Permutations for E*cologists. R package version 1.1.4. Available online at: https://CRAN.R-project.org/package=asnipe

Faulkes, C. G., Bennett, N. C., Bruford, M. W., O’brien H. P., Aguilar G. H., & Jarvis, J. U. M. (1997). Ecological constraints drive social evolution in the African mole–rats. Proceedings of the Royal Society of London B: Biological Sciences, 267. doi:10.1098/rspb.1997.0226

Fox, J., & Weisberg, S. (2011). An {R} Companion to Applied Regression, Second Edition. Thousand Oaks CA: Sage. URL: http://socserv.socsci.mcmaster.ca/jfox/Books/Companion

Getz, L. L., & Hofmann, J. E. (1986). Social organization in free-living prairie voles, *Microtus ochrogaster*. Behavioral Ecology and Sociobiology, 18, 275–282.

Getz, L. L., Hofmann, J. E., McGuire, B., & Dolan, T. W. III. (2001). Twenty-five years of population fluctuations of *Microtus ochrogaster* and *M. pennsylvanicus* in three habitats in east-central Illinois. Journal of Mammalogy, 82, 22–34.

Getz, L. L., & McGuire, B. (1993). A comparison of living singly and in male-female pairs in the prairie vole, *Microtus ochrogaster*. Ethology, 94, 265–278. doi:10.1111/j.1439-0310.1993.tb00444.x

Getz, L. L., McGuire, B., Pizzuto, T., Hofmann, J. E., & Frase, B. (1993). Social organization of the prairie vole (*Microtus ochrogaster*). Journal of Mammalogy. 74, 44–58. doi: 10.2307/1381904

Getz, L. L., McGuire, B., Hofmann, J., Pizzuto, T., & Frase, B. (1994). Natal dispersal and philopatry in prairie voles (*Microtus ochrogaster*): settlement, survival, and potential reproductive success. Ethology Ecology & Evolution, 6, 267–284.

Hatchwell, B. J., & Komdeur, J. (2000). Ecological constraints, life history traits and the evolution of cooperative breeding. Animal Behaviour, 59, 1079–1086. doi:10.1006/anbe.2000.1394

Jetz, W., & Rubenstein, D. R. (2011). Environmental uncertainty and the global biogeography of cooperative breeding in birds. Current Biology, 21, 72–78. doi:10.1016/j.cub.2010.11.075.

Kappeler, P. M., Cremer, S., & Nunn, C. L. (2015). Sociality and health: impacts of sociality on disease susceptibility and transmission in animal and human societies. Philosophical Transactions of the Royal Society B: Biological Sciences, 370. doi:10.1098/rstb.2014.0116

Keane, B., Bryant, L., Goyal, U., Williams, S., Kortering, S. L., Lucia, K. E., Richmond, A. R., & Solomon, N. G. (2007). No effect of body condition at weaning on survival and reproduction in prairie voles. Canadian Journal of Zoology, 85, 718–727.

Kleiber, C., & Zeileis, A. (2008). Applied econometrics with R. New York: Springer-Verlag. ISBN 978-0-387-77316-2. https://CRAN.R-project.org/package=AER” https://CRAN.R-project.org/package=AER

Krause, J., & Ruxton, G. D. (2002). Living in groups. Oxford: Oxford University Press.

Lambert, C. T., Sabol, A. C., & Solomon, N. G. (2018). Genetic monogamy in socially monogamous mammals is primarily predicted by multiple life history factors: a meta-analysis. Frontiers in Ecology and Evolution, 6, 139. doi:10.3389/fevo.2018.00139

Langwig, K. E., Frick, W. F., Bried, J. T., Hicks, A. C., Kunz, T. H. & Marm Kilpatrick, A. (2012), Sociality, density-dependence and microclimates determine the persistence of populations suffering from a novel fungal disease, white-nose syndrome. Ecology Letters, 15: 1050–1057. doi:10.1111/j.1461-0248.2012.01829.x

Lemen, C. A., & Freeman, P. W. (1985). Tracking mammals with fluorescent pigments: a new technique. Journal of Mammalogy, 66, 134–136.

Lott, D. F. (1991). Intraspecific variation in the social systems of wild vertebrates. Cambridge: Cambridge University Press.

Lucia, K. E., Keane, B., Hayes, L. D., Lin, Y. K., Schaefer, R. L., & Solomon, N. G. (2008). Philopatry in prairie voles: an evaluation of the habitat saturation hypothesis. Behavioral. Ecology, 19, 774–783. doi: 10.1093/beheco/arn028

Lukas, D., & Clutton-Brock, T. H. (2013). The evolution of social monogamy in mammals. Science, 341, 526–530. doi:10.1126/science.1238677

Lukas, D., & Clutton-Brock, T. H. (2017). Climate and the distribution of cooperative breeding in mammals. Royal Society Open Science, 4. doi:10.1098/rsos.160897

Lukas, D., & Huchard, E. (2014). The evolution of infanticide by males in mammalian societies. Science, 346, 841–844. doi:0.1126/science.1257226

Lutermann, H., Bennett, N. C., Speakman, J. R., & Scantlebury, M. (2013). Energetic benefits of sociality offset the costs of parasitism in a cooperative mammal. PLOS ONE, 8, e57969. doi:10.1371/journal.pone.0057969

Mabry, K. E., Streatfeild, C. A., Keane, B., & Solomon, N. G. (2011). avpr1a length polymorphism is not associated with either social or genetic monogamy in free-living prairie voles. Animal Behaviour, 81, 11–18.

McGuire, B., & Getz, L.L. (2010). Alternative male reproductive tactics in a natural population of prairie voles, *Microtus ochrogaster*. Acta Theriol, 55, 261–270. doi:10.4098/j.at.0001-7051.077.2009

McGuire, B., Getz, L. L., & Oli, M. K. (2002). Fitness consequences of sociality in prairie voles, Microtus ochrogaster: influence of group size and composition. Animal Behaviour, 64, 645–654. doi:10.1006/anbe.2002.3094.

McGuire, B., Pizzuto, T., Getz, L.L. (1990). Potential for social interaction in a natural population of prairie voles (*Microtus ochrogaster*). Canadian Journal of Zoology, 68, 391–398.

Nunn, C.L., Jordán, F., McCabe, C.M., Verdolin, J.L., Fewell, J.H. (2015). Infectious disease and group size: more than just a numbers game. Philosophical Transactions of the Royal Society B, 370, 20140111.

Okhovat, M., Berrio, A., Wallace, G., Ophir, A. G., & Phelps, S. M. (2015). Sexual fidelity trade-offs promote regulatory variation in the prairie vole brain. Science, 350, 1371–1374.

Ophir, A.G., Wolff, J.O., Phelps, S.M. (2008). Variation in neural V1aR predicts sexual fidelity and space use among male prairie voles in semi-natural settings. PNAS, 105, 1249–1254.

Psorakis, I., Roberts, S. J., Rezek, I., & Sheldon, B. C. (2012). Inferring social network structure in ecological systems from spatio-temporal data streams. Journal of the Royal Society Interface, 9, 3055–3066. doi: 10.1098/rsif.2012.0223

R Core Team (2017). R: A Language and Environment for Statistical Computing. Vienna: R Foundation for Statistical Computing. Available online at: https://www.R-project.org/.

Richmond, M., & Conaway, C. H. (1969). Management, breeding, and reproductive performance of the vole, *Microtus ochrogaster*, in a laboratory colony. Laboratory Animal Care, 19, 80–87.

Rubenstein, D. R., & Lovette, I. J. (2007) Temporal environmental variability drives the evolution of cooperative breeding in birds. Current Biology, 17, 1414–1419. doi:10.1016/j.cub.2007.07.032.

Ryder, T.B., Parker, P.G., Blake, J.G., & Loiselle, B.A. (2009). It takes two to tango: reproductive skew and social correlates of male mating success in a lek-breeding bird. Proceedings of the Royal Society B: Biological Sciences, 276, 2377–2384. doi:10.1098/rspb.2009.0208

Sabol, A. C., Solomon, N. G., & Dantzer, B. (2018). How to study socially monogamous behavior in secretive animals? Using social network analyses and automated tracking systems to study the social behavior of prairie voles. Frontiers in Ecology and Evolution, 6, 178. doi:10.3389/fevo.2018.00178

Schradin, C., & Pillay, N. (2005). Intraspecific variation in the spatial and social organization of the African striped mouse, Journal of Mammalogy, 86, 99–107.

Schradin, C., König, B., Pillay, N. (2010). Reproductive competition favours solitary living while ecological constraints impose group-living in African striped mice. Journal of Animal Ecology, 73, 515–521.

Schülke, O., Bhagavatula, J., Vigilant, L., & Ostner, J. (2010). Social bonds enhance reproductive success in male macaques. Current Biology, 20, 2207–2210. doi:10.1016/j.cub.2010.10.058.

Shen, S-F., Emlen, S.T., Koenig, W.D., Rubenstein, D.R. (2017). The ecology of cooperative breeding behaviour. Ecology Letters, 20, 708–720.

Shuster, S. M., & Wade, M. J. (2003). Mating systems and mating strategies. Princeton: Princeton University Press.

Shuster, S. M., Willen, R. M., Keane, B., & Solomon, N. G. (2019). Alternative mating tactics in socially monogamous prairie voles, Microtus ochrogaster. Frontiers in Ecology and Evolution, 7. doi:10.3389/fevo.2019.00007

Sikes, R. S., Bryan, J. A., Byman, D., Danielson, B. J., Eggleston, J., Gannon, M. R., Gannon, W. L., Hale, D. W., Jesmer, B. R., Odell, D. K., Olson, L. E., Stevens, R. D., Thompson, T. A., Timm, R. M., Trewhitt, S. A., & Willoughby, J. R. (2016). 2016 Guidelines of the American Society of Mammalogists for the use of wild mammals in research and education. Journal of Mammalogy 97, 663–688.

Silk, J. B. (2007). The adaptive value of sociality in mammalian groups. Philosophical Transactions of the Royal Society B: Biological Sciences, 362, 539–559. doi:10.1098/rstb.2006.1994

Silk, J. B., Alberts, S. C., & Altmann, J. (2003). Social bonds of female baboons enhance infant survival. Science, 302, 1231–1234. doi:10.1126/science.1088580

Silk, J.B., Beehner, J.C., Bergman, T.J., Crockford, C., Engh, A.L., Moscovice, L.R., Wittig, R.M., Seyfarth, R.M., Cheney, D.L. (2009). The benefits of social capital: close social bonds among female baboons enhance offspring survival. Proceedings of the Royal Society B, 276, 3099–3104.

Silk, J.B., Beehner, J.C., Bergman, T.J., Crockford, C., Engh, A.L., Moscovice, L.R., Wittig, R.M., Seyfarth, R.M., Cheney, D.L. (2010). Strong and consistent social bonds enhance the longevity of female baboons. Current Biology, 20, 1359–1361.

Solomon, N. G. (1993). Comparison of parental behavior in male and female prairie voles (*Microtus ochrogaster*). Canadian Journal of Zoology, 71, 1991–1994.

Solomon, N. G., & Keane, B. (2018). Dispatches from the field: sociality and reproductive success in prairie voles. Animal Behaviour, 143, 193–203. doi:10.1016/j.anbehav.2018.07.001

Solomon, N. G., & Jacquot, J. J. (2002). Characteristics of resident and wandering prairie voles, *Microtus ochrogaster*. Canadian Journal of Zoology, 80, 951–955.

Solomon, N. G., Keane, B., Knoch, L. R., & Hogan, P. J. (2004). Multiple paternity in socially monogamous prairie voles (Microtus ochrogaster). Canadian Journal of Zoology, 82, 1667–1671.

Solomon, N.G., Richmond, A. R., Harding, P. A., Fries, A., Jacquemin, S., Schaefer, R. L., Lucia, K. E., & Keane, B. (2009). Polymorphism at the avpr1a locus in male prairie voles correlated with genetic but not social monogamy in field populations. Molecular Ecology, 18, 4680–4695.

Spiegel, O., Sih, A., Leu, S. T., & Bull, C. M. (2017). Where should we meet? Mapping social network interactions of sleepy lizards shows sex-dependent social network structure. Animal Behaviour, 136, 207–215. doi:10.1016/j.anbehav.2017.11.001.

Streatfeild, C. A., Mabry, K. E., Keane, B., Crist, T. O., & Solomon, N. G. (2011). Intraspecific variability in the social and genetic mating systems of prairie voles, *Microtus ochrogaster*. Animal Behaviour, 82, 1387–1398. doi: 10.1016/j.anbehav.2011.09.023

van Schaik, C. (1983). Why are diurnal primates living in groups? Behaviour, 87, 120–144.

Walum, H., & Young, L. J. (2018). The neural mechanisms and circuitry of the pair bond. Nature Reviews Neuroscience, 19, 643–654. doi:10.1038/s41583-018-0072-6

Whitehead, H. (2008). Analyzing animal societies: quantitative methods for vertebrate social analysis. Chicago: University of Chicago Press.

Whiteman, N. K., & Parker, P. G. (2004). Effects of host sociality on ectoparasite population biology. Journal of Parasitology, 90, 939–947. doi:10.1645/GE-310R

Wickham, H. (2009). Ggplot2: Elegant graphics for data analysis. New York: Springer-Verlag

Young, L. J., & Wang, Z. (2004). The neurobiology of pair bonding. Nature Neuroscience, 7, 1048–1054. doi:10.1038/nn1327

